# Stress granule formation enables anchorage-independence survival in cancer cells

**DOI:** 10.1101/2024.09.14.613064

**Authors:** Seungwon Yang, Anaïs Aulas, Paul J. Anderson, Pavel Ivanov

## Abstract

Stress granules (SGs) are dynamic cytoplasmic structures assembled in response to various stress stimuli that enhance cell survival under adverse environmental conditions. Here we show that SGs contribute to breast cancer progression by enhancing the survival of cells subjected to anoikis stress. SG assembly is triggered by inhibition of Focal Adhesion Kinase (FAK) or loss of adhesion signals. Combined proteomic analysis and functional studies reveal that SG formation enhances cancer cell proliferation, resistance to metabolic stress, anoikis resistance, and migration. Importantly, inhibiting SG formation promotes the sensitivity of cancer cells to FAK inhibitors being developed as cancer therapeutics. Furthermore, we identify the Rho-ROCK- PERK-eIF2α axis as a critical signaling pathway activated by loss of adhesion signals and inhibition of the FAK-mTOR-eIF4F complex in breast cancer cells. By triggering SG assembly and AKT activation in response to anoikis stress, PERK functions as an oncoprotein in breast cancer cells. Overall, our study highlights the significance of SG formation in breast cancer progression and suggests that therapeutic inhibition of SG assembly may reverse anoikis resistance in treatment-resistant cancers such as triple-negative breast cancer (TNBC).

**Highlights:**

- Either anoikis stress or loss of adhesion induce stress granule (SG) formation
- The Rho-ROCK-PERK-eIF2α axis is a crucial signaling pathway triggered by the absence of adhesion signals, leading to the promotion of SG formation along with the inhibition of the FAK- AKT/mTOR-eIF4F complex under anoikis stress.
- PERK functions as an oncogene in breast cancer cells, initiating SG formation and activating AKT under anoikis stress.
- Inhibiting SG formation significantly enhances the sensitivity to Focal Adhesion Kinase (FAK) inhibitors, suggesting a potential for combined therapy to improve cancer treatment efficacy.

## Introduction

Stress granules (SGs) are non-membranous cytoplasmic condensates that contain translationally-stalled mRNAs, associated translation initiation factors and multiple RNA-binding proteins (RBPs)[1–4]. These structures are dynamically assembled in response to adverse environmental conditions including heat shock, oxidative stress, osmotic stress, UV damage, and viral infection, and disassemble when stress is relieved [4–8]. Cells that are genetically resistant to SG assembly (e.g., G3BP1/2^-/-^) are more vulnerable to oxidative (e.g., sodium arsenite) stress-induced death [9–11], findings that have identified SGs as components of the integrated stress response (ISR) pathway that protects cells exposed to adverse environmental conditions.

SGs of distinct compositions and functions are assembled in response to different stresses. “Canonical” SG assembly is initiated by phosphorylation of eIF2α by Heme-regulated initiation factor 2 α kinase (HRI, eIF2αK1), Protein kinase RNA-activated (PKR, eIF2αK2), PKR-like endoplasmic reticulum kinase (PERK, eIF2αK3), or General control nonderepressible 2 (GCN2, eIF2αK4) resulting in reduced translation initiation. This pathway disables the eIF2 ternary complex, resulting in stalled translation and assembly of SGs. “Non-canonical” SG assembly is initiated by stresses that inhibit the mammalian target of rapamycin (mTOR) kinase. Downregulated mTOR activity results in the dephosphorylation of 4EBP1 which inhibits the assembly of eIF4F, to inhibit translation initiation and induce SG assembly [3, 12–14].

The propensity for cancer cells to metastasize correlates with the development of anchorage independence and resistance to anoikis, the tendency for cells removed from a substrate to undergo programmed cell death. These properties facilitate survival in the circulatory system and invasion of distant tissues [15, 16]. Several signaling molecules that promote SG assembly are also linked to anoikis, suggesting a functional link between these processes: i) Loss of cell adhesion, an upstream trigger of anoikis, inhibits AKT, the kinase that activates mTOR to inhibit translation initiation and induce SG assembly [17–19], ii)Rho GTPases play a prominent role in the interplay between cellular adhesion and anoikis. Specifically, RhoA-ROCK signaling induces both SG assembly, and anoikis [17, 20], iii) PERK, the eIF2a kinase that induces SG assembly is an oncoprotein that is activated by ROCK [21–23]. These correlations suggested that SGs may contribute to cancer metastasis by promoting resistance to anoikis.

Here we report a previously unappreciated link between SGs and anoikis. We show that the loss of adhesion signals that trigger anoikis also induces SG assembly in response to PERK- eIF2α phosphorylation and functional inactivation of the eIF4F complex to allow survival of detached cells. This pathway is likely responsible for the chemosensitization of triple-negative breast cancer (TNBC) cells by treatments that inhibit SG assembly [24]. This suggests that inhibition of SG assembly could act as an adjuvant therapy in the treatment of TNBC, even in cases where treatment is extremely challenging due to the lack of specific targets unlike other subtypes of breast cancers

## 1. Stress granule proteins are associated with cancer progression

Although several studies have reported that overexpression of certain SG proteins can enhance cancer progression, the role of SGs, per se, in this process has not been determined [25–29]. To address this issue, we expressed wild type and mutant G3BP1 (G3BP1-ΔNTF2, lacking the G3BP dimerization domain) in G3BP1/2^-/-^ cells and quantified SA-induced SG assembly [Fig.1A]. The G3BP1/2^-/-^ cells transfected with G3BP1-ΔNTF2 completely block stress granule (SG) formation [30]. The immunoprecipitation (IP) samples were quality-controlled using well-known stress granule components such as eIF4G, USP10, and FXR1 [Fig.1B].

**Figure 1.**
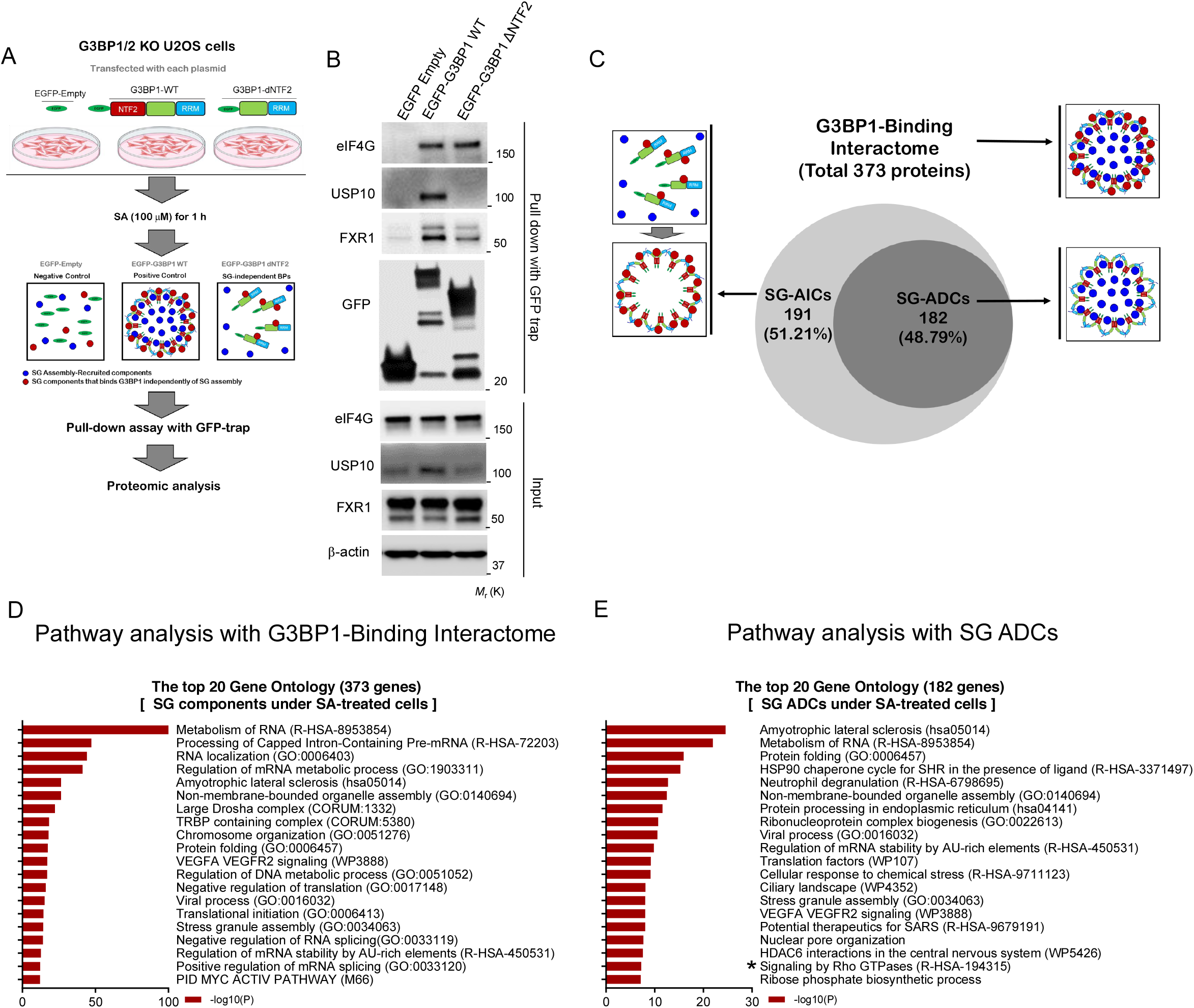
Stress granule-associated component is associated with cancer progression. A. Schematic summary how to identify SA-induced SG components. B. Pulldown assay showing the quality control of each binding partners for EGFP-empty, EGFP- G3BP1 WT, and EGFP-G3BP1 ΔNTF2 in wellknown SG components. C. Quantitative Venn diagram showing the number of overlapping and non-overlapping SG components as assembly-dependent and assembly-independent components, respectively. D. Top 20 enriched pathways up-regulated by G3BP1-Binding Interactome under SA. E. Top 20 enriched pathways up-regulated by SG assembly dependent component (In the G3BP1-Binding Interactome, components that were bound to the G3BP1 mutant (ΔNTF2) were eliminated to identify SG assembly-dependent components under SA.)

Our G3BP1-interactome analysis identified a total of 373 proteins of which 191 bound G3BP1 constitutively (assembly-independent components, SG-AICs), whereas 182 were newly recruited during SA-induced SG assembly (assembly-dependent components, SG-ADCs) [Fig.1C and Supplementary Fig. S1A and B]. Our pathway analysis identified components of the TRBP complex, VEGF signaling, HDAC6 interaction, and Signaling by Rho GTPase pathway [Fig.1D and E]. These signaling pathways are closely associated with driving cancer development and progression [31–34]. Specifically, the Rho GTPase pathways directly control numerous factors associated with cancer behaviors and progression, including adhesion, migration, invasion, anoikis resistance, epithelial-mesenchymal transition, and mechanotransduction [35]. Taken together, our results indicate SG assembly has a potential for regulating cancer activities and progression.

## 2. Stress granule positively controls cancer cell activities

To assess the role of SGs in cancer cell functions such as proliferation, resistance to metabolic stress, anoikis resistance, and migration, we compared these functional responses in cells with greater or lesser propensity for SG assembly. The functional responses of cells lacking G3BP [Supplementary Fig. S2A-S2C], or G3BP1/2^-/-^ cells reconstituted with wild-type G3BP1 or G3BP1ΔNTF2 were found to differ significantly. U2OS (G3BP1/2^-/-^) cells lacking the capacity to assemble SGs exhibit reduced proliferation [Fig 2A-D], increased sensitivity to metabolic stress [Fig. 2E-H] increased susceptibility to anoikis [Fig. 2I-L], and reduced migration [Fig. 2M-R] compared to WT controls. These effects were reversed when G3BP1/2^-/-^ cells were reconstituted with WT G3BP1, but not with the G3BP1 (ΔNTF2) mutant that is incapable of conferring the ability to assemble SGs [Figs. C, D, K, L, N, O, Q, R). Thus, cells that are unable to assemble SGs are deficient in cell proliferation, migration, and survival under adverse conditions, all of which are critical properties of cancer cells.

**Figure 2.**
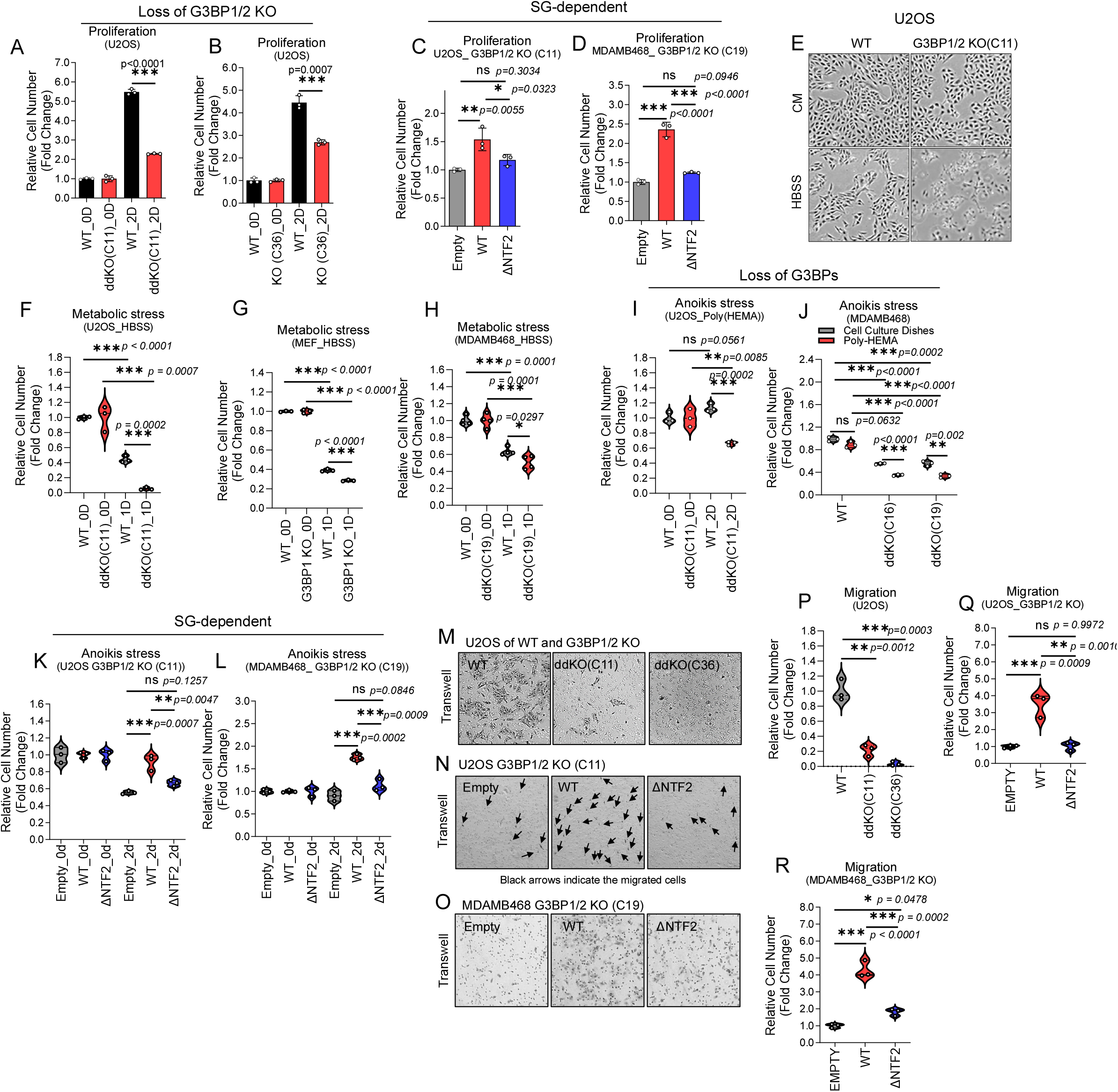
Stress granule up-regulates cancer cell activities. A and B. Cell proliferation analysis in WT and G3BP1/2^-/-^ cells of U2OS cells cultured with 0.5% FBS cell culture medium for 2 days (2D). C and D. Cell proliferation analysis in G3BP1/2^-/-^ cells with EGFP-empty, EGFP-G3BP1 WT, or EGFP-G3BP1 ΔNTF2. U2OS and MDAMB468 cells were cultured with 0.5% and 10% FBS for 2 days, respectively. E. Representative images for WT and G3BP1/2^-/-^ cells with or without HBSS for 24h. F-H. Analysis of resistance to metabolic stress in WT, G3BP1/2^-/-^ or ΔG3BP1 of U2OS, MEF, and MDAMB468 for 1 day (1D). I-L. Analysis for resistance to anoikis stress in WT and G3BP1/2^-/-^, or G3BP1/2^-/-^ cells transfected with EGFP-empty, EGFP-G3BP1 WT, or EGFP-G3BP1 ΔNTF2 in U2OS and MDAMB468. M-O. Representative images of cell migration in P-R, P-R. Cell migration assay in WT and G3BP1/2^-/-^, or G3BP1/2^-/-^ transfected with EGFP-empty, EGFP-G3BP1 WT, or EGFP-G3BP1 ΔNTF2 in U2OS and MDAMB468. Statistical analysis: Student’s t-test (A-B and F-J), Anova (C-D, K-L, and Q-R), Mean ± SD are shown. *, p< 0.05; **, p< 0.01; ***, p< 0.001.

## 3. Adhesion signals inversely correlate with eIF2α phosphorylation

In our SG interactome analysis, SG dynamics have been closely associated with the Rho GTPase pathway. Rho GTPases are widely recognized as pivotal regulators of intracellular actin dynamics, which contribute to multiple cellular processes such as cell mitosis, cell adhesion, and cell migration [18, 36–38]. Rho GTPases, particularly Rho and downstream targets, are temporally activated to induce mitotic cell rounding and cortical stiffening during the ingression phase of cytokinesis during mitosis [36, 39, 40]. Upon cell detachment, Rho and downstream targets are quickly upregulated to induce actomyosin cortical actin to produce tension under the cell membrane, allowing the cell to adopt a spherical shape [17, 41]. In addition, local activation or inactivation of individual Rho GTPases including Rho, Rac, and CDC42, is also required for polarized cell movements produced by coordinate spatio-temporal cytoskeletal dynamics [42, 43]. For example, during cell migration, Rho activation mainly promotes detachment of the trailing edge of a moving cell via actomyosin contraction, whereas activation of Rac and CDC42 drive cell migration via lamellipodium and filopodium extension, respectively at the leading edge of mobile cells [44]. Given that Rho GTPase is implicated in the processes of cell attachment and detachment[17, 45, 46], and considering that Rho GTPase signaling is coupled with SG assembly-dependent components, our findings suggest a hypothesis whereby SGs may play a role in stress conditions related to cell adhesion signaling. These associations led us to focus on understanding the physiological function of SGs under anoikis stress.

We first assessed whether inhibiting adhesion signals by a specific inhibitor initiates SG-related signal pathways. The inhibition of Focal Adhesion Kinase (FAK), a central mediator of adhesion signals, leads to the phosphorylation of eIF2α and the dephosphorylation of 4EBP1, both of which trigger SG assembly. The phosphorylation levels of FAK, Paxillin and AKT are inversely correlated with adhesion strength. Inhibition of FAK suppresses these adhesion signals in a dose-dependent manner in U2OS cells [Fig.3A], as well as breast cancer cell lines including SKbr3, MCF7, MDAMB361, and MDAMB231 [Fig.3B-E].

**Figure 3.**
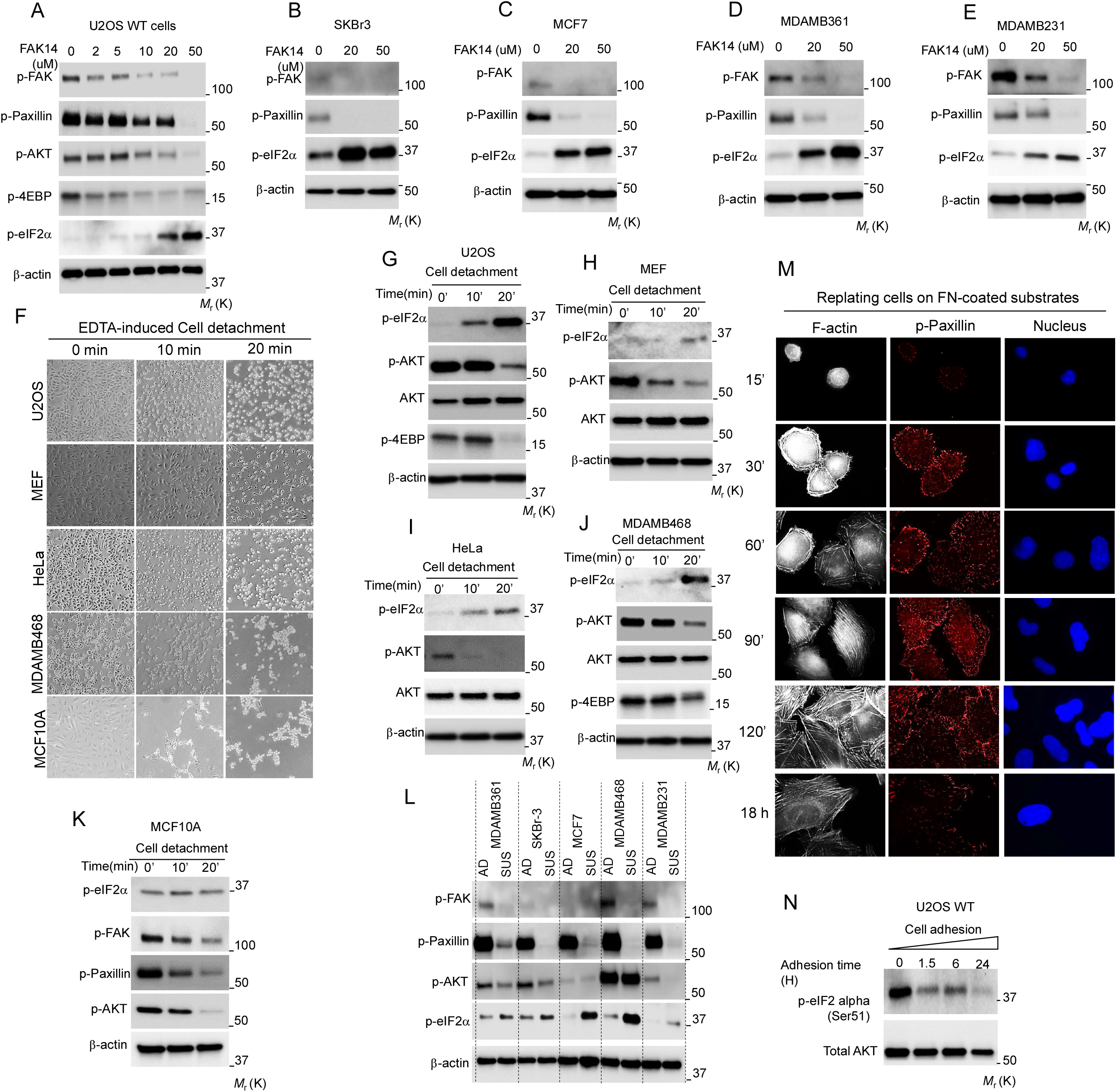
Adhesion signals reversely regulate eIF2α phosphorylation. A. Immunoblot analysis of the protein levels under FAK14 treatment for 1h in U2OS cells. B-E. Immunoblot analysis of the protein levels under FAK14 treatment for 1h in SKbr3, MCF7, MDAMB361, or MDAMB231. p-FAK and p-Paxillin were used as a marker of adhesion signals. F. Representative images for EDTA-induced cell detachment. EDTA (Final conc. 5mM) was added into cell culture medium to dynamically induce cell detachment. G-K. Immunoblot analysis of the protein levels under EDTA (Final conc. 5mM)-induced cell detachment in U2OS, MEF, HeLa, MDAMB468, and MCF10A. L. Immunoblot analysis of the protein levels under EDTA (Final conc. 5mM)-induced cell detachment for 20min in MDAMB361, SKBr3, MCF7, MDAMB468, and MDAMB231 M. Representative serial IF images during cell adhesion. N. Immunoblot analysis of the protein levels during cell adhesion.

Next, we determined whether physical and dynamic cell detachment also induces eIF2α phosphorylation and 4EBP1 dephosphorylation. EDTA was directly added to cell culture medium to induce cell detachment without cleavage of membrane receptors including adhesion molecules and growth receptors unlike treatment with trypsin [47]. EDTA was found to induce phosphorylation of eIF2α and cellular detachment with similar kinetics in U2OS, MEF, HeLa, MDAMB468, MCF10A, MDAMD361, SKBr3, MCF7, and MDAMB231 cell lines [Fig.4F-L].

**Figure 4.**
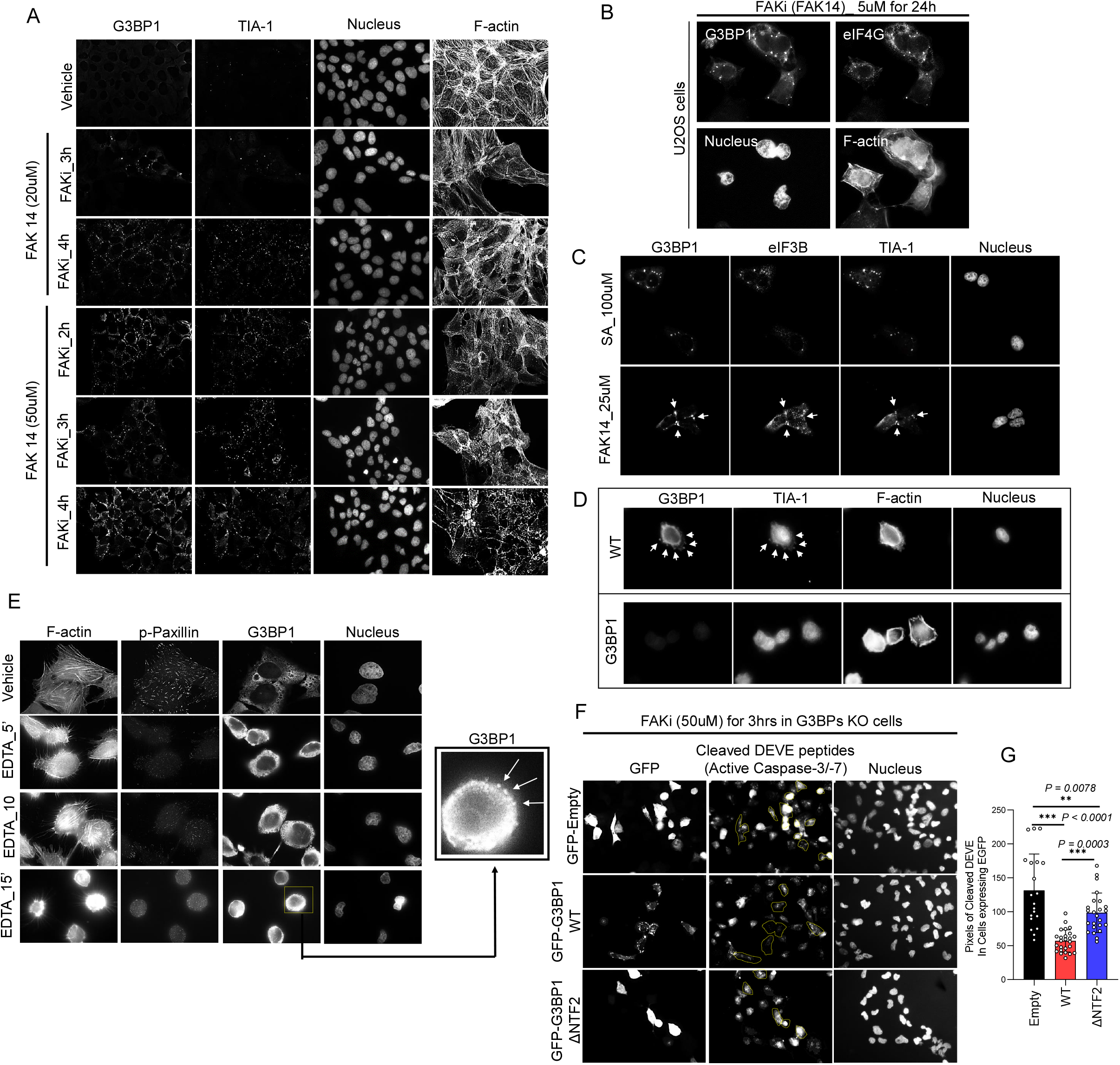
Adhesion signals reversely regulate SG formation. A. Analysis of stress granule formation induced by FAKi in a dose- and time-dependent manner. B. Analysis of stress granule formation induced by low dose of FAKi for 24h. C. Analysis of canonical stress granule stained with eIF3B and TIA-1 under SA (100uM, 1h) or FAKi (25uM, 4h) treatment. D. Analysis of stress granule formation in WT and G3BP1/2^-/-^ U2OS cells during cell adhesion. Cells were fixed 15min after seeded on fibronectin-coated slide glasses to visualize SGs at the early of adhesion stage. White arrows indicate SGs. E. Analysis of stress granule formation in EDTA (2mM)-induced cell detachment, F-actin and p-Paxillin were used as a marker of adhesion signals and cell morphology. White arrows indicate SG formation. F. Representative images for SG formation and active caspase-3 and -7 in G3BP1/2^-/-^ U2OS cells transfected with EGFP-empty, EGFP-G3BP1 WT, or EGFP-G3BP1 ΔNTF2. Yellow dots indicate the transfected cells with EGFP-tagged each proteins. Caspase activities were detected by CellEvent™ Caspase-3/7 Detection Reagents (C10423). G. Quantification of apoptotic cells stained with active Caspase-3 and -7. Statistical analysis: Anova (G) *, p< 0.05; **, p< 0.01; ***, p< 0.001.

Interestingly, normal cell lines such as MEFs and MCF10A cells exhibit relatively less eIF2α phosphorylation in response to anoikis stress than cancer cell lines [Fig.3H and K]. In addition, increased cell adhesion is inversely correlated with eIF2α phosphorylation [Fig. 3M and N]. These results indicate that anoikis stress dynamically induces eIF2α phosphorylation and 4EBP1dephosphorylation to enhance SG formation.

## 4. FAK inhibition induces canonical stress granule formation

Consistent with our conjecture that loss of cell adhesion signals can trigger SG assembly, the FAK inhibitor (FAKi) induces SG assembly in a dose- and time-dependent manner [Fig.4A and Supplementary Fig. S3A]. Exposure to low dose FAK inhibitor for prolonged periods also induces SG assembly [Fig.4B]. These are “canonical” pro-survival SGs as judged by their inclusion of eIF3B, G3BP1 and TIA-1[48, 49] [Fig.4C]. Importantly, FAK inhibition gradually induced cell detachment without causing complete cell death. This is likely due to the disruption of inside-out signaling through the FAK–integrins axis, resulting in alterations in cell morphology [50].

Although the small spherical morphology of cells in suspension precluded visualization of SGs, we were able to observe SGs when these cells weakly attached to substrates during the early stages of cell adhesion, when anoikis stress was still present. WT and G3BP1/2^-/-^ cells were cultured in suspension for 1 hour, then seeded onto fibronectin-coated glass and fixed after 15 minutes for SG analysis. SGs were observed in WT, but not G3BP1/2^-/-^ cells at the initial attachment stage [Fig.4D]. Conversely, we observed SGs in cells that had almost completely detached in response to EDTA treatment [Fig.4E and Supplementary Fig. S3B].

To determine whether SG assembly could increase resistance to anoikis stress, EGFP-Empty, EGFP-G3BP1, and EGFP-G3BP1 ΔNTF2 were transfected into G3BP1/2^-/-^ cells prior to induction of FAK i-induced anoikis stress. Only cells expressing EGFP-G3BP1 formed SGs. In these cells FAK inhibitor induced significantly lower levels of caspase-3 and -7 activity compared to EGFP-empty or EGFP-G3BP1 ΔNTF2 controls [Fig.4F and G]. Taken together, our findings demonstrate that canonical SG formation is induced by loss of adhesion signals and is correlated with resistance to anoikis stress.

## 5. Loss of adhesion signaling induces SGs including unique SG components through both ROCK-PERK activation and mTORC-eIF4F complex inhibition

To identify the eIF2α kinase responsible for anoikis stress-induced phosphorylation of eIF2α, we compared the phosphorylation of signaling molecules in adherent and EDTA-induced suspension cells lacking GCN2, PKR, HRI, and PERK [51]. ΔPERK cells uniquely failed to phosphorylate eIF2α and inhibit protein synthesis in response to anoikis stress [Fig.5A and B, Supplementary Fig. S4A and S4B]. PERK KO cells were also impaired in their ability to induce SG assembly in response to FAKi [Fig.5C and D]. We conclude that PERK is the eIF2α kinase that acts upstream of anoikis stress. PERK is a downstream target for the Rho-associated protein kinase (ROCK) [21] that enhances SG formation [20]. RhoA-ROCK is strongly activated by loss of adhesion signaling [17, 52] and is known to enhance the survival of cancer cells exposed to anoikis stress [46]. Taken together, ROCK may participate as an upstream signal in a PERK-eIF2α-SG axis. Consistent with this hypothesis, ROCK inhibitors significantly downregulate FAKi-induced SG assembly [Fig.5E and F]. As inhibition of either PERK or ROCK was unable to completely inhibit FAKi-induced SG formation, an additional signaling pathway may be involved in anoikis stress-induced SG assembly. The ability of FAKi to promote hypophosphorylation of 4EBP1 [Fig.5G] suggests that inhibition of the eIF4F complex could also contribute to FAKi-induced SG assembly. To test this hypothesis, the eIF4F complex was isolated using m^7^GTP-Sepharose under various stress conditions, including treatment with SA, FAKi, or mTORi. FAKi, like mTORi, disrupted the eIF4F complex [Fig. 5G], suggesting that anoikis induces SG formation through both the PERK and mTOR pathways.

**Figure 5.**
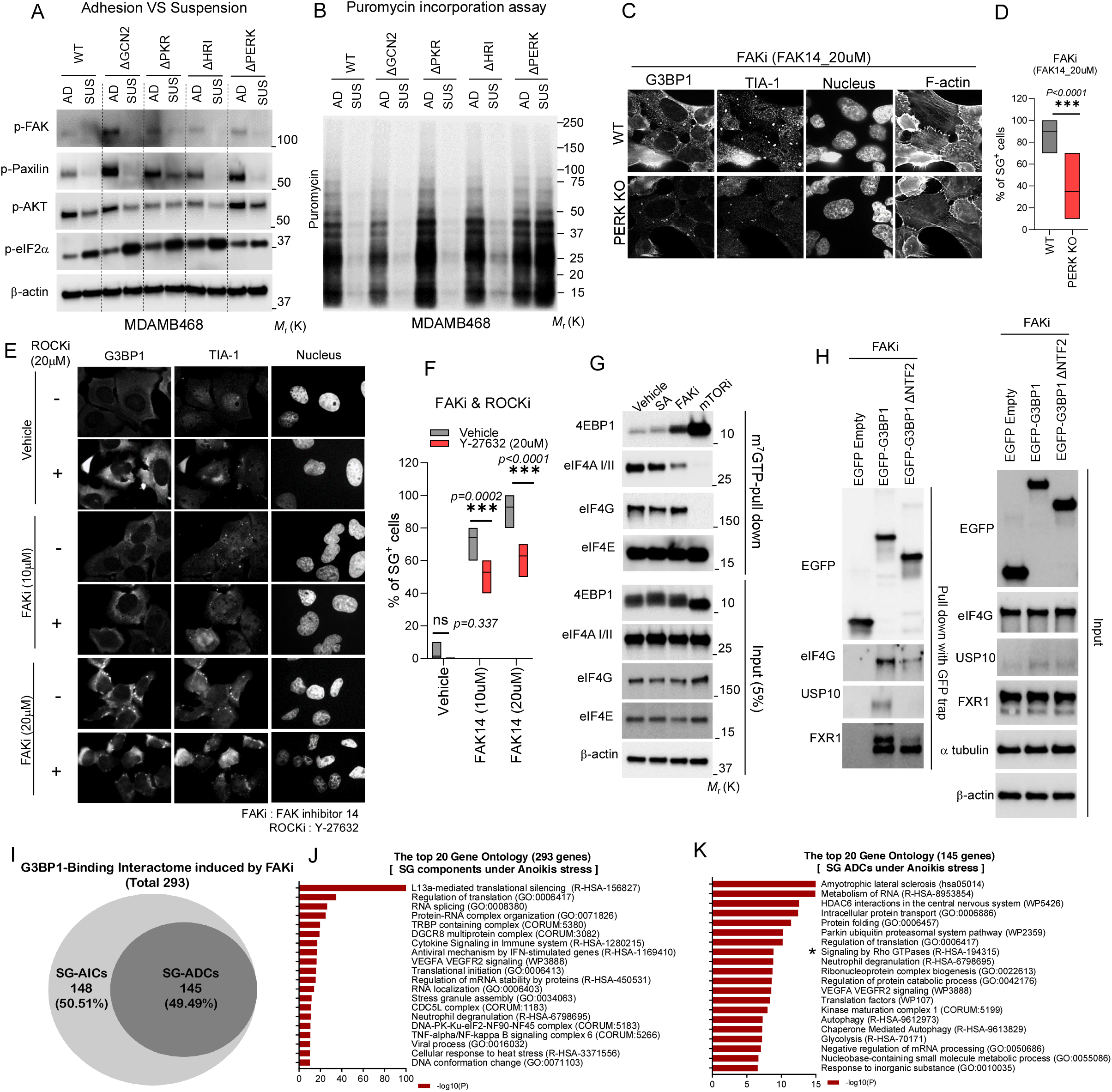
Both ROCK-PERK activation and mTORC-eIF4F complex inhibition induces SG formation under anoikis stress. A. Immunoblot analysis of the protein levels with or without EDTA (2mM, 15min)-induced cell detachment in WT, ΔGCN2, ΔPKR, ΔHRI, and ΔPERK of MDAMB468 (AD: Adhesion, and SUS: Suspesion). B. Puromycin incorporation assay under EDTA (2mM, 15min)-induced cell detachment in WT,ΔGCN2, ΔPKR, ΔHRI, and ΔPERK of MDAMB468. C. Representative images for WT and ΔPERK of Hap1 cells under FAKi-induced SG formation. D. Quantification of FAKi-induced SG formation in WT and ΔPERK cells. E. Representative images for FAKi-induced SG formation with or without ROCKi treatment. F. Quantification of FAKi-induced SG formation with or without ROCKi treatment. G. m^7^GTP pulldown assay under treatment of SA (100uM), FAK14 (50uM), or Torin-1 (10uM) for 2h in U2OS cells. H. Pulldown assay for EGFP-empty, EGFP-G3BP1 WT, and EGFP-G3BP1 ΔNTF2 showing partners (well known SG components, the quality control for binding). I. Quantitative Venn diagram showing the number of overlapping and non-overlapping SG components as assembly-dependent components (ADC) and assembly-independent components (AIC), respectively. J. Top 20 enriched pathways up-regulated by G3BP1-Binding Interactome under FAKi. K. Top 20 enriched pathways up-regulated by SG assembly dependent component (In the G3BP1-Binding Interactome, components that were bound to the G3BP1 mutant (ΔNTF2) were eliminated to identify SG assembly-dependent components under FAKi).

Importantly, previous research has demonstrated stress-specific differences in the assembly and composition of stress granules (SGs)[53]. To elucidate the differences in SG components formed under anoikis stress compared to those induced by SA, we performed a proteomic analysis using the methodology outlined in Figure 1 [Fig 5H and I]. In comparing the G3BP1- Binding Interactome, we identified that among the 293 FAKi-induced SG components, 103 were unique. Furthermore, when focusing SG assembly-dependent components, only 42 out of 145 (28.9%) overlapped with SA-induced SG components, whereas 103 components (71%) were unique to FAKi-induced SGs [Supplementary Fig. S5A and B]. Our pathway analysis reveals SG components linked to signaling by RhoGTPase is more enriched in FAKi-induced SG components than in SA-induced SG components [Fig. 5J and K, and Supplementary Fig. S5C and D]. Intriguingly, the cancer-associated pathways related to Programmed cell death, Kinase maturation complex, and Autophagy are top-ranked among the FAKi-induced SG components [Fig. 5K, Supplementary Fig. S5D][54–56]. These results suggest that anoikis-induced SG may contribute more to cancer activities compared to SGs formed by other types of stress.

## 6. CASP4 is one of SG downstream targets in Anoikis-induced SG assembly

Based on our proteomic analysis, we have surmised that programmed cell death signaling is implicated in anoikis-induced SG assembly. These SGs can sequester pro-apoptotic factors, including caspase-3 and caspase-7, thereby inhibiting apoptosis during SG formation[9]. Interestingly, our interactome analysis revealed that anoikis-induced SGs do not contain caspase-3 or -7 typically present in SA-induced SGs [Supplementary Fig. 5A, B, and D]. Notably, CASP4 (caspase-4) was the only caspase identified in the anoikis-induced SG-associated ADCs [Supplementary Fig. 5A (A13 in FAKi-induced unique G3BP1 interactome), B (A13 in FAKi-induced unique SG ADCs), and D]. CASP4 is recognized as an inflammatory caspase that activates the non-canonical inflammasome, leading to pyroptosis through the activation of gasdermin D (GSDMD)[57]. Furthermore, activated GSDMD can also trigger the activation of caspase-3 to induce cell death[58]. Indeed, CASP4 is present in FAKi-induced SGs [Fig. 6A], and CASP4 is consistently upregulated and hyperactivated by FAKi in G3BP1/2^-/-^ U2OS cells compared to WT U2OS cells [Fig. 6B and Supplementary Fig. 6A].

**Figure 6.**
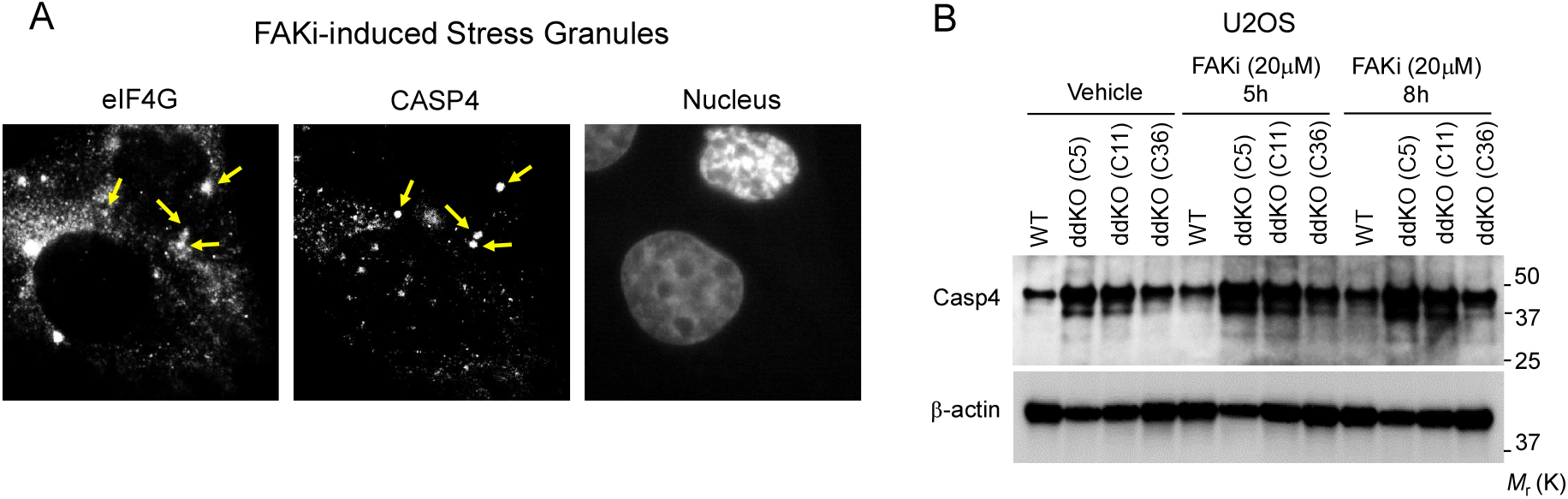
CASP4 is one of SG downstream targets in Anoikis-induced SG assembly. A. Analysis of stress granule formation induced by FAK14 (20μM) for 4h in U2OS cells, and Yellow arrows indicate co-localization between eIF4G and CASP4. B. Immunoblot analysis of the protein levels under FAK14 treatment in WT and G3BP1/2^-/-^ of U2OS cells.

## 7. PERK is an oncoprotein that induces anoikis resistance in breast cancer cells

We found that eIF2α phosphorylation in response to anoikis stress is much lower in non-cancerous breast cells compared to breast cancer cells [Fig.3K], and that PERK contributes to anoikis-induced eIF2α phosphorylation and SG assembly [Fig.5A and B]. PERK mRNA are overexpressed in various cancer patient tissues such as Acute Myeloid Leukemia (AML), Breast cancer, Esophageal Cancer, Lung Adenocarcinoma (AC), Lung Squamous Cell Carcinoma (SC), Pancreatic Cancer, Prostate Cancer, Rectal Cancer, Renal Cell Carcinoma (CC), Renal Chromophobe Carcinoma (CH), Renal Papillary Carcinoma (PA), Testicular Cancer, and Uterine Endometrial Carcinoma (EC). [Fig.7A and Supplementary Fig. 7A], and PERK protein is highly upregulated in most subtypes of breast cancer cell lines, including ER-positive, HER2- positive, and Triple-Negative Breast Cancer (TNBC) [Fig.7B]. In addition, absence of PERK reduced proliferation and anoikis resistance of a TNBC cell line under both normal and PolyHEMA substrate respectively [Fig.7C]. Moreover, PERK has been shown to activate key survival signals in cancer, including AKT, mTOR, and ERK [22, 23]. We therefore determined whether PERK expression enhances the activation of AKT and mTOR in breast cancer cells.. Treatment with FAKi or EDTA-induced anoikis stress rapidly reduced AKT and mTOR activity, which are dependent on adhesion signaling, during the early stages of anoikis stress [Fig. 3A-K]. However, PERK expression remains high in breast cancer cells [Fig.7B]. Therefore, overexpressed PERK might promote survival under stress conditions, in part via anoikis-induced SG formation. We hypothesized that under sustained anoikis conditions, the significantly increased ROCK activity would activate PERK, consequently leading to the restoration of survival signals, including AKT, that were quickly downregulated due to loss of adhesion signals at the early stage of anoikis stress [Fig. 3A-K]. To conduct these experiments, cells were detached using EDTA to minimize membrane receptor damage. Cells in suspension were replated on dishes coated with Poly(HEMA) [Fig. 7D]. Consistent with our hypothesis, MDAMB468 breast cancer cells showed higher AKT and mTOR activity under anoikis stress than non-cancerous breast cells [Fig.7E]. Moreover, TNBCs lacking PERK were not able to maintain high AKT activity under sustained anoikis conditions. PERK in cancer cells also seemed to play a role in restoring AKT activity under anoikis conditions [Fig.7F]. These results suggest that PERK in breast cancer cells induces SG formation at an early stage of anoikis stress and also preserves AKT activity during sustained stress conditions to support SG assembly and cell survival.

**Figure 7.**
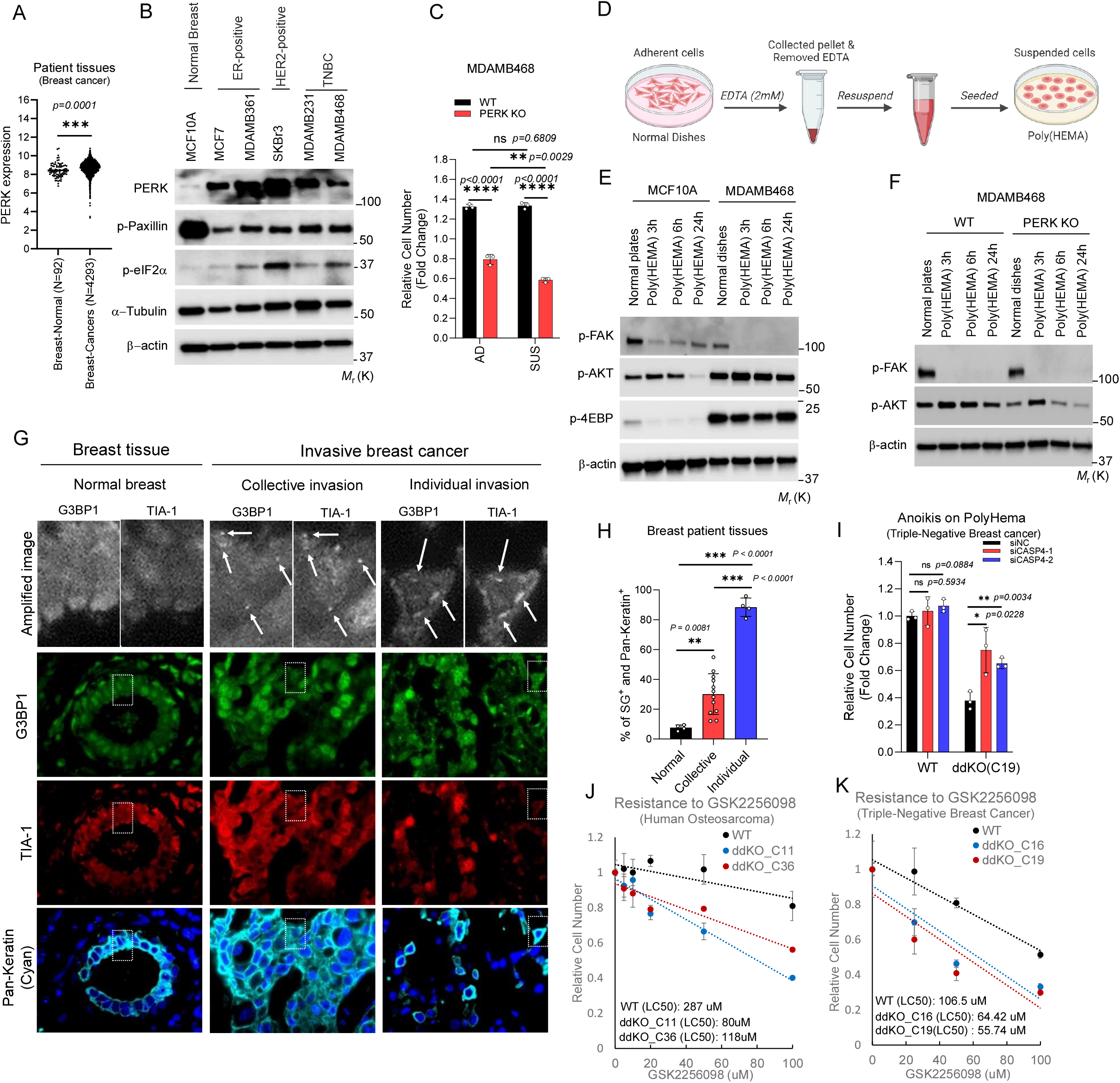
PERK is required for the upregulated resistance to anoikis stress in MDAMB468 cells. A. PERK expression between normal and breast cancer patient tissues derived from pubulic data set of GENT2 (http://gent2.appex.kr/gent2/). B. Analysis of PERK expression in various breast cancer cell lines. p-Paxillin was used for basal adhesion signals, and α-tubulin and β-actin were used as loading controls. C. Proliferation assay in WT and ΔPERK of MDAMB468 with or without cell adhesion. D. Schematic summary how to track AKT activity during long term anoikis stress. E. Immunoblot analysis of p-AKT and p-4EBP levels under long term anoikis stress in nomal breast cells(MCF10A) and TNBC cells (MDAMB468). F. Immunoblot analysis of p-AKT levels under long term anoikis stress in WT and ΔPERK of MDAMB468. G. Representative IF staining images of each tissue group. White arrows inducate colocalization of G3BP1 and TIA-1 as typical SGs (Tissue Mictoarray_BR087e) H. Quantification of SG-positive cells in normal, collective invasion, or individual invasion tissues. I. Analysis for resistance to anoikis stress in WT and G3BP1/2^-/-^ of MDAMB468 with siNC or siCASP4. J. Analysis of resistance to FAKi (GSK2256098) in WT and G3BP1/2^-/-^ of U2OS cells. K. Analysis of resistance to FAKi (GSK2256098) in WT and G3BP1/2^-/-^ of MDAMB468 TNBC cells. Statistical analysis: Statistical analysis: Student’s t-test and Anova, *, p< 0.05; **, p< 0.01; ***, p< 0.001.

## 8. Inhibiting SG formation potentiates the anti-cancer activity of FAKi

We discovered that FAK inhibition or anoikis stress leads to the formation of SGs. These SGs play a crucial role in conferring resistance to anoikis stress, and we applied this knowledge to analysis of human breast cancer tissue and anticancer agents targeting FAK.

Metastasis-initiating cancer cells often undergo a loss of firm attachment to the extracellular matrix (ECM) in their primary tissue. These metastatic cells are continuously exposed to weak adhesion signals due to their poorly attached states. Particularly in breast cancer patient tissue, the individual invasive cancer cells encounter relatively weaker adhesion signals, with RhoGTPase playing a critical role[59, 60]. Therefore, we investigated the presence of SGs in patient tissues of breast cancer cells. Consistent with our conjecture, SGs were observed in collective invading cancer cells and nearly all individual invading cancer cells, which are highly challenging for cancer detection. [Fig.7G and H, and Supplementary Fig.6A and B]. Furthermore, loss of CASP4 significantly rescued the defect of anoikis resistance in G3BP1/2^-/-^ MDAMB468 cells [Fig. 7I and Supplementary Fig. 7D].

FAK has been identified as a promising anticancer target, and several FAK inhibitors (FAKis) are currently undergoing clinical trials [61–64]. We found that the antiproliferative activity of the FAKi GSK2256098 is potentiated in U2OS and TNBC cells that lack G3BP1/2 and are therefore unable to assemble SGs [Fig.7J and K].

These results suggest that SGs are indeed present in metastatic and invading cancer cells in human breast cancer patient tissue, and combining inhibition of SG formation with FAK inhibitor could provide a more effective treatment strategy. This approach enables the reduction of FAK inhibitor (FAKi) doses, thereby minimizing potential side effects associated with high concentrations of FAKi.

## Discussion

In this study, we have elucidated a previously unrecognized function of SGs in enhancing resistance to anoikis stress in breast cancer cells, thus promoting cancer progression and metastasis [Supplementary Fig.7E].

Our findings provide reliable evidence that SG formation is a critical adaptive response that enables cancer cells to survive under conditions of inappropriate adhesion, a characteristic feature of metastatic dissemination.

We first showed that cell adhesion signals are inversely correlated with SG assembly. Both anoikis stress (loss of adhesion) and Focal Adhesion Kinase (FAK) inhibition can induce SG formation. This new finding represents a significant advancement in the SG field, suggesting that SGs are not only a response to traditional stressors such as oxidative stress, heat shock, and UV damage response but also play a critical role in cellular adaptation to adhesion-linked stresses. We further identified the Rho-ROCK-PERK-eIF2α and the FAK-AKT/mTOR-eIF4F complex axes, which are activated and inhibited by the loss of adhesion signals as a pivotal signaling pathways, leading to the promotion of SGs under anoikis stress. The dual regulatory pathways enhance anchorage-independence survival in metastatic cancer cells. PERK, a well-known eIF2α kinase, has been shown to function as a protumorigenic factor in breast cancer cells. Our results reveal that PERK not only initiates SG formation but also maintains AKT activity during prolonged anoikis stress. This maintenance of AKT activity is essential for cell survival and highlights the multifaceted role of PERK in promoting cancer cell adaptability and metastatic potential. One of the most significant findings of our study is that inhibiting SG formation markedly enhances the sensitivity of cancer cells to FAK inhibitors. This suggests a promising therapeutic strategy where the combination of SG inhibitors and FAK inhibitors could be used to effectively target and eliminate cancer cells, particularly in treatment-resistant cancers such as triple-negative breast cancer.

Furthermore, our proteomic analysis, based on the GFP-Trap with nanobody (to overcome the high background issues commonly associated with conventional antibody-based approaches) and G3BP1 ΔNTF2 mutant for inhibiting G3BP dimerization reveals SG assembly-recruited components is more linked in cancer progression-associated pathways including signaling by RhoGTPase, which is critical for cell proliferation, migration, and invasion in cancer metastatic environment. In addition to previous proteomic analysis that analyzed SG components with or without stress, we distinguished between SG assembly-dependent and independent components by using G3BP1 ΔNTF2 only to inhibit SG formation under the same stress condition. The newly recruited SG components during SG assembly, identified as SG assembly-dependent components, were more enriched in pathways related to cancer progression.

Among anoikis-induced SG components, we first identified CASP4 as a unique SG assembly-dependent component for programed cell death, which is the highly ranked SG-associated pathway, in the context of anoikis stress. We found that the inhibition of CASP4 by SG assembly contributes to enhanced anoikis resistance in breast cancer cells. The identification of CASP4 as a key player in anoikis resistance highlights the potential of targeting SG-related mechanisms as a therapeutic approach in anoikis resistant cancers.

Interestingly, the anoikis-induced SG mechanism incorporates both major SG formation pathways: eIF2α phosphorylation and inhibition of the mTOR-eIF4F complex. Previous studies have linked eIF2α to canonical SG formation and cell survival, while inhibition of the mTOR- eIF4F complex has been associated with non-canonical SGs and cell death. Canonical SGs have been shown to sequester pro-apoptotic factors, thereby impeding apoptosis and promoting cell survival. In contrast, non-canonical SGs, lacking these apoptotic factors, have been suggested to enhance programmed cell death [48, 49, 65, 66]. However, this binary understanding may not be always absolute. For instance, non-canonical SGs induced by UV exposure, nitric oxide exposure or H_2_O_2_ can sometimes promote cell survival [7, 49, 53, 67–69]. Additionally, our findings indicate that SGs can confer resistance to metabolic stress, a condition known to induce non-canonical SGs [65].

Given these observations, it is evident that the functional outcomes of SGs are not solely determined by their canonical or non-canonical nature. Transcriptomic and proteomic alterations, which can vary depending on cell type and stress conditions, may significantly influence the balance between pro-survival and pro-death signals within SGs. As a result, the function of SGs can vary widely depending on the cellular context, making it difficult to generalize their roles beyond cytoprotection.

In addition, our live-cell imaging unveiled a dynamic equilibrium between SG assembly and disassembly, constantly assembling and disassembling, in response to anoikis. As stress gradually accumulated, the rate of SG disassembly decreased, resulting in an upward trend in SG assembly. Based on these observations, we hypothesize that large SGs may form under conditions of severe stress, while tiny SGs may assemble and disassemble dynamically during routine cellular processes. These findings suggest that SGs may play a role not only in response to acute stress but also in more delicate cellular processes such as mitosis, differentiation, and cytokine responses. Further investigation is warranted to fully elucidate the role of SGs in these processes.

In summary, this study uncovers a novel function of stress granules (SGs) in promoting anoikis resistance and metastasis in breast cancer cells. We demonstrate that loss of adhesion induces SG formation via activating the Rho-ROCK-PERK-eIF2α and inhibiting FAK-AKT/mTOR-eIF4F pathways, highlighting SGs as critical players in cellular adaptation to adhesion-linked stresses. Our findings reveal that PERK is pivotal not only for initiating SG formation but also for maintaining AKT activity under prolonged anoikis stress, essential for cell survival.

Furthermore, inhibiting SG formation sensitizes cancer cells to FAK inhibitors, offering a promising therapeutic approach for aggressive cancers like triple-negative breast cancer (TNBC). Advanced proteomic analyses identify SG assembly-dependent components, such as CASP4, that are linked to cancer progression and anoikis resistance. Our study emphasizes the significance of SGs in cancer biology and suggests that targeting SG-related pathways could enhance the efficacy of existing cancer therapies.

### Materials and methods Cell culture and Cell lines

U2OS, MEF, MDAMB468, SKBr, MCF7, MDAMB361, MDAMB231, HeLa, MCF10A, and Hap1 were maintained at 37 °C in a 5% CO2 incubator, and cultured in DMEM (Corning) supplemented with10% FBS, 100 U/ml penicillin, and 100 µg/ml streptomycin. The cells and KO cell lines such as U2OS-WT & ΔΔG3BP1/2 cells, MEF-WT & ΔG3BP1 cell, MDAMB468-WT, ΔΔG3BP1/2, ΔGCN2, ΔPKR, ΔHRI & ΔPERK, and Hap1-WT & ΔPERK cells were established and used for previous studies[13, 70].

## 7. Methyl GTP Sepharose Chromatography

eIF4F complex was captured by m^7^GTP-Sepharose bead as described previously[71], In brief, m^7^GTP-Sepharose bead was washed twice with ice-cold cell 1X IP buffer (Cell Signaling, #9803) including 20 mM Tris-HCl (pH 7.5), 150 mM NaCl, 1 mM Na_2_EDTA, 1 mM EGTA, 1% Triton, 2.5 mM sodium pyrophosphate, 1 mM beta-glycerophosphate, 1 mM Na_3_VO_4_, 1 µg/ml leupeptin, and 1 mM PMSF. U2OS cells cultured with the indicated inhibitors and harvested with IP buffer. Each cell supernant lysate was incubated with pre-washed m^7^GTP-Sepharose, and incubated at 4°C for 1 hour with shaking on a rotator. Unbound proteins were removed by washing 3 times with the IP Buffer, and proteins bound to m^7^GTP-Sepharose were eluted with SDS loading buffer.

### Immunoblotting

Immunoblotting was peformed as described previously[17, 72]. Cells were washed twice with ice-cold 1X PBS including 1 mM Sodium Fluoride, 1 mM Na_3_VO_4_, and 1 mM PMSF to remove residual FBS, to prevent dephosphorylation of phosphotated proteins, and to inhibit protein degradation during cell pellet washing and harvest for attached or suspended cells. The pre-washed cells was lysed with RIPA buffer (Cell Signaling, #9806) containing 20 mM Tris-HCl (pH 7.5), 150 mM NaCl, 1 mM Na_2_EDTA, 1 mM EGTA, 1% NP-40, 1% sodium deoxycholate, 2.5 mM sodium pyrophosphate, 1 mM beta-glycerophosphate, 1 mM Na_3_VO4, 1 µg/ml leupeptin, and 1 mM PMSF. Cell lysates were resolved by SDS-PAGE and transferred onto nitrocellulose membranes followed by blotting.

### GFP Trap pull-down assay with Plasmid DNA transfection

U2OS ΔΔG3BP1/2 cells were transfected with EGFP-Empty, EGFP-G3BP1, and G3BP1- ΔNTF2 plasmids using transfection reagent using GenJet™ Plus DNA In Vitro Transfection Reagent (Signagen, # SL100499) for 2 days. Cells were washed twice with ice-cold 1X PBS, and lysed with 1X IP buffer (Cell Signaling, #9803). ChromoTek GFP-Trap® Magnetic Agarose bead(Proteintech, # gtma) was washed twice with ice-cold cell 1X IP buffer (Cell Signaling, #9803) by Magnetic stand. Each cell supernant lysate was incubated with the pre-washed GFP-Trap® Magnetic bead at 4°C for 1 hour with shaking on a rotator. Unbound proteins were removed by washing 3 times with the IP Buffer, and proteins bound to GFP-Trap® Magnetic bead were eluted with SDS loading buffer. The analysis of binding partners of G3BP1 was performed by Taplin Mass Spectrometry Facility in Harvard Medical School.

### siRNA transfection

Cells were transfected with siRNA targeting negative control (siNC_Ambion, Invitrogen, Cat#:4390843, Lot# ASO2MT2K), or an individual gene (siCASP4-1_Ambion, Invitrogen, Ref: 4427038 (CASP4), LOT:ASO2MYN1, ID s2414, or siCASP4-2 Ambion, Invitrogen, Ref: 4427038 (CASP4), LOT:ASO2MYN2, ID s2413). siRNAs were transfected using PepMute™ siRNA Transfection Reagent (Signagen, Ijamsville, MD), according to the manufacturer’s instructions.

### Pathway and Venn diagram analysis

Gene ontology (GO) enrichment and Kyoto Encyclopedia of Genes and Genomes (KEGG) pathway enrichment analyses were performed using Metascape (https://metascape.org/), a web-based portal designed to provide a comprehensive gene list annotation and analysis resource, as described previously[73, 74]. Quantitative Venn diagrams were generated using the following web tool: https://www.deepvenn.com/

### Migration assay

Migration assays were performed as described previously[72]. Briefly, the cells were suspended in 150 µl of serum-free medium and seeded onto 8-mm Pore Transwell Inserts (Corning, #353097) for the migration assay. The lower chamber was filled with 900 µl of 10% FBS complete medium. After 24hr, the cells on the Transwell Inserts were then fixed with 4% paraformaldehyde/PBS for 30 min. Subsequently, fixed cells were stained with hematoxylin solution (Sigma-Aldrich, #MHS16) for 2h. After wiping off cells on the upper side of the filter on the Transwell Inserts using cotton swabs, microphotographs of the cells migrated onto the lower side of the filter were taken using light microscopy. Cells migrated onto the lower side of the filter were manually counted from the microphotographs. Mean cell numbers were quantified from randomly selected same size of squares per Transwell insert.

### Anoikis assay

Model substrate for anoikis assay was used as described previously[17, 75]. Briefly, A solution of 10 mg/ml poly-HEMA (Sigma-Aldrich, #P3932) mixed in 95% ethanol was added onto cell culture dishes and dried in the tissue culture hood. Once completely dried, the dishes were washed with PBS extensively at least 2 times. After incubating cells in poly(HEMA)-coated dishes, Cell Counting Kit (CCK8) (Abcam, #ab228554) was directly added into the poly(HEMA)- coated dish with or without suspended cells as a negative control to measure relative cell number following the manufacturer’s protocol, as described previously[17, 72]. The final CCK-8 values were normalized by the CCK-8 values obtained from early seeded cells as the starting point CCK-8 values.

### Assays for Cell Proliferation and cell viability under metabolic stress

Cell proliferatation and metbolic stress assay were performed using Cell Counting Kit (CCK8) (Abcam, #ab228554) as described previously[17, 72]. Cells were seeded for 1 day to firmly attach the cells on cell culture dishes. The attached cells were washed twice with Hank’s Balanced Salt Solution with calcium and magnesium (HBSS) and replaced with the indicated culture buffers such as 10% FBS, 0.5% FBS, and HBSS with calcium and magnesium. CCK-8 reaction buffer was prepared in 5% FBS DMEM, and was used to measure CCK-8 values. The final CCK-8 values were normalized by the CCK-8 values obtained from early seeded cells as the starting point CCK-8 values.

### Human Breast Tumor Samples and IHC images

Tissue Microarray for Human breast and breast cancer tissue was obtained from TissueArray (TISSUEARRAY, # BR087e). Representative H&E staining image for the IF staining was obtained from Tissue assay (BR087e) in the website (tissuearray.com/tissue-arrays/Breast/BR087e).

### Ribopuromycylation

Ribopuromycylation was performed as described previously[13]. Briefly, 5 minutes prior to end of cell harvest treatment, puromycin (final concentration of 9 µM) was add directly to cell culture dishes. The treated cells were incubated for 5 min at 37°C. The cells were harvested and prepared for standard immunoblotting. Puromycin incorporation was detected by anti-puromycin antibody (Millipore, #MABE343 clone 12D10, IgG2a) for immunoblotting.

### Immunofluorescence (IF) of cells and tissue

Immunofluorescence analysis was performed as described previously[13, 17, 72, 76]. For immunofluorescence analysis of cells, cells on slide glass were fixed with 4% paraformaldehyde/PBS for 30 min and permeabilized with 1% Triton X-100 (Sigma-Aldrich, T8787)/PBS for 20 min. Cells were then washed with PBS with 0.1% Triton-X100 (IF washing buffer). The samples were then incubated with a blocking solution of 1% albumin from chicken egg white (Sigma-Aldrich, A5503) in the IF washing buffer for 1 h. The cells were incubated with primary antibodies in the IF washing buffer for 1 h, then the cells were washed 3 times for 10 min with the IF washing buffer. The cells was incubated with secondary antibodies and Rhodamine Phalloidin Reagent or Phalloidin FITC Reagent (Abcam, # ab235138 or #ab235137) in the IF washing buffer for 1 h, then the cells were washed 3 times for 10 min with the IF washing buffer. Hoechst 33258 (Sigma-Aldrich, # H21486) was added during the second washing step.

For immunofluorescence analysis of tissues, formalin-fixed, paraffin-embedded tissue sections, The tissue slides were deparaffinized and rehydrated. Briefly, tissue slides were placed on a slide warmer at 62°C for 20 min, and melted paraffin carefully wiped off to prevent tissue detachment. The tissue slides were immersed in 100% Xylene three times, each time for 3 minutes. Then, the tissue slides were immersed in 100% ethanol three times, each time for 3 minutes to completely remove residual Xylene. Serially, the slides were immersed in 95% ethanol for 3 min, 70% ethanol for 3 min, 50% ethanol for 3 min, and rinsed in water for 5 min. For antigen retrieval, the slides were incubated in Tris-EDTA buffer (consisting of 1.27 ml of 1M EDTA solution, 5 ml of 2M Tris-HCl, and 0.5 ml of Tween 20 per 1 liter of H2O, with pH 7.5) at 90°C to 95°C for 20 min. The slides were rinsed in cold tap water for 10 min. The slides were then incubated with a blocking solution of 1% albumin from chicken egg white (Sigma-Aldrich, A5503) in the IF washing buffer for 1 h. The tissue were incubated with primary antibodies in the IF washing buffer for 1 h, then washed 3 times for 10 min with the IF washing buffer. The slides were incubated with secondary antibodies in the IF washing buffer for 1 h, then washed 3 times for 10 min with the IF washing buffer. Hoechst 33258 (Sigma-Aldrich, # H21486) was added during the second washing step.

## Acknowledgments

We deeply appreciate Yuichiro Adachi, Claire Riggs, and Prakash Kharel for their technical expertise, comprehensive knowledge, and protocol support, along with their assistance in data analysis. Additionally, we are grateful to Victoria Ivanova, Allison Williams, Nupur Bhatter, and Safiyah Zubair for supplying experimental reagents, offering technical insights and support. Lastly, we extend our thanks to Shawn M. Lyons for his valuable research feedback. We also acknowledge the Brigham and Women’s Hospital Confocal Microscopy Core for their support with live imaging.

This work was supported by funds from National Institutes of Health grant R35 GM126901 (P.J.A.), National Institutes of Health grant R01 GM126150 and R01 GM146997 (P.I.),

---

**Supplementary Figure S1, Related figure 1.**
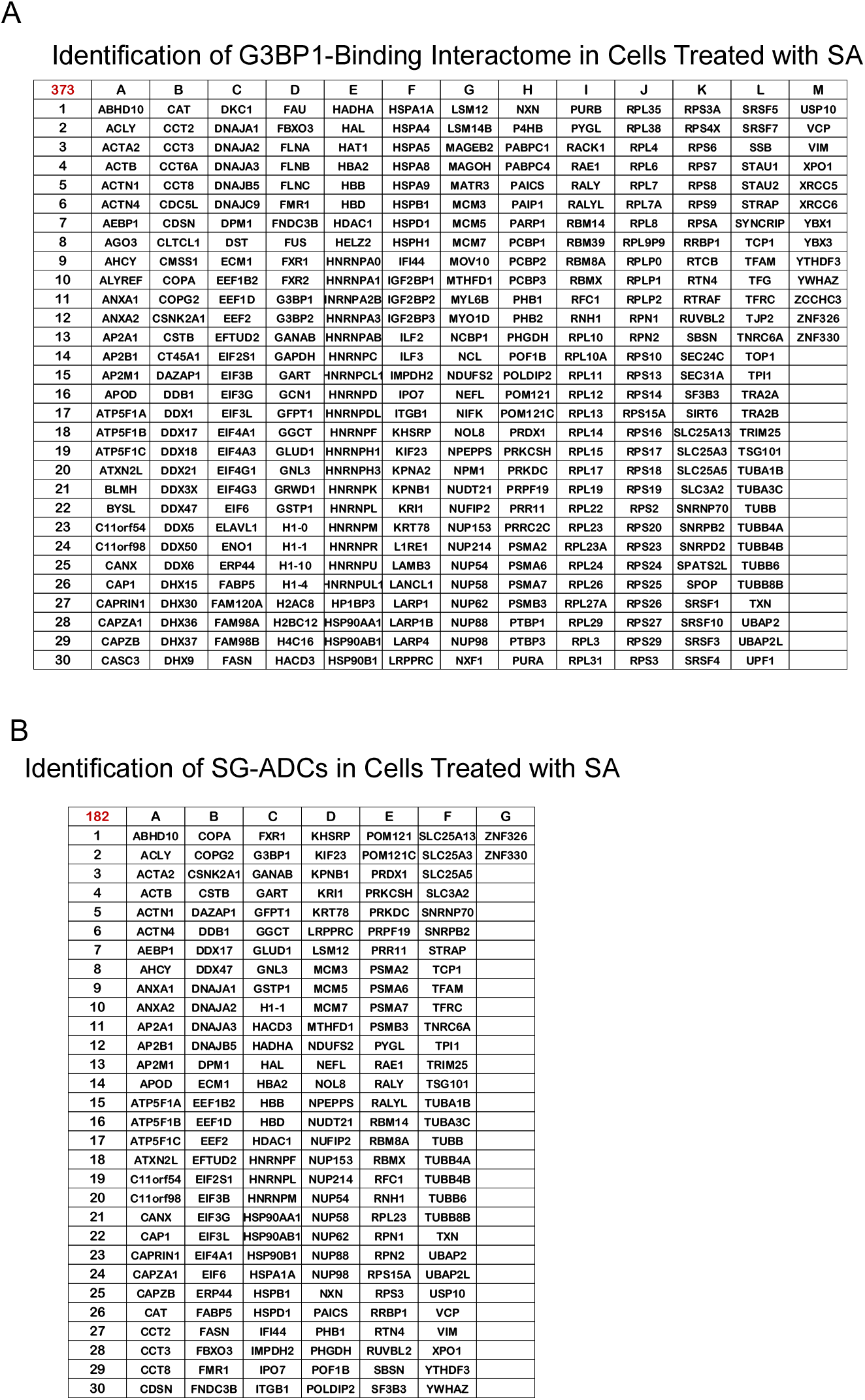
A. Identification of G3BP1-Binding Interactome in U2OS cells treated with SA. B. Identification of SG-ADCs in U2OS cells treated with SA.

**Supplementary Figure S2, Related figure 2.**
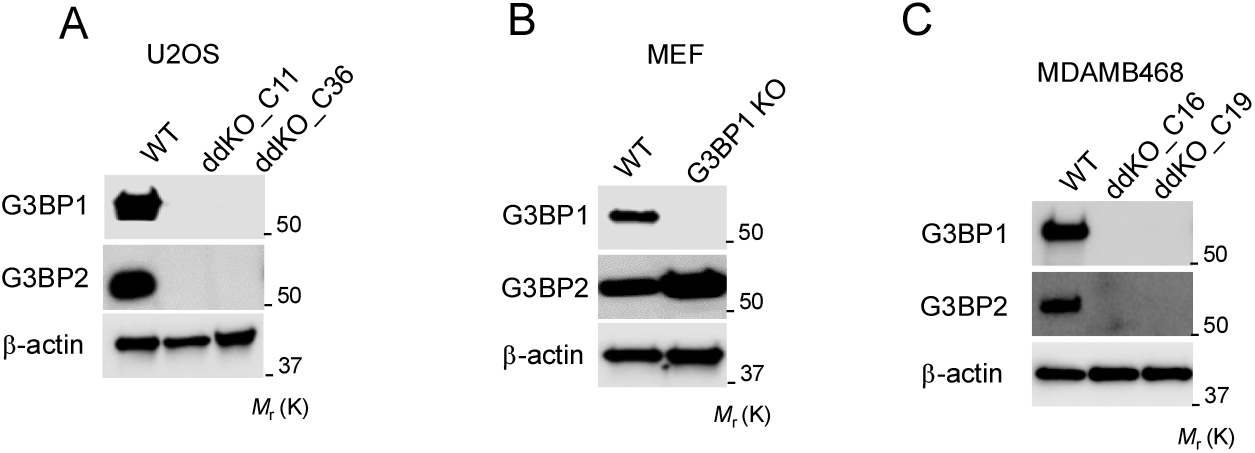
A-C. An immunoblot analysis of the G3BP1 or 2 protein levels in different KO cell clones.

**Supplementary Figure S3, Related figure 4.**
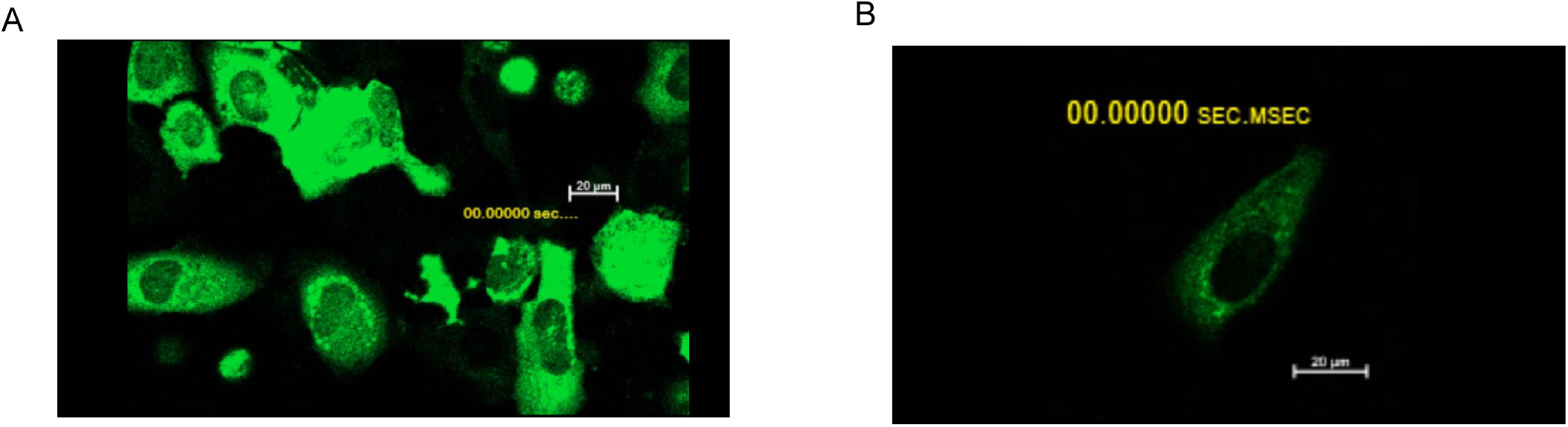
A. Live cell image for FAKi-induced SG assembly B. Live cell image for EDTA-induced SG assembly

**Supplementary Figure S4, Related figure 5.**
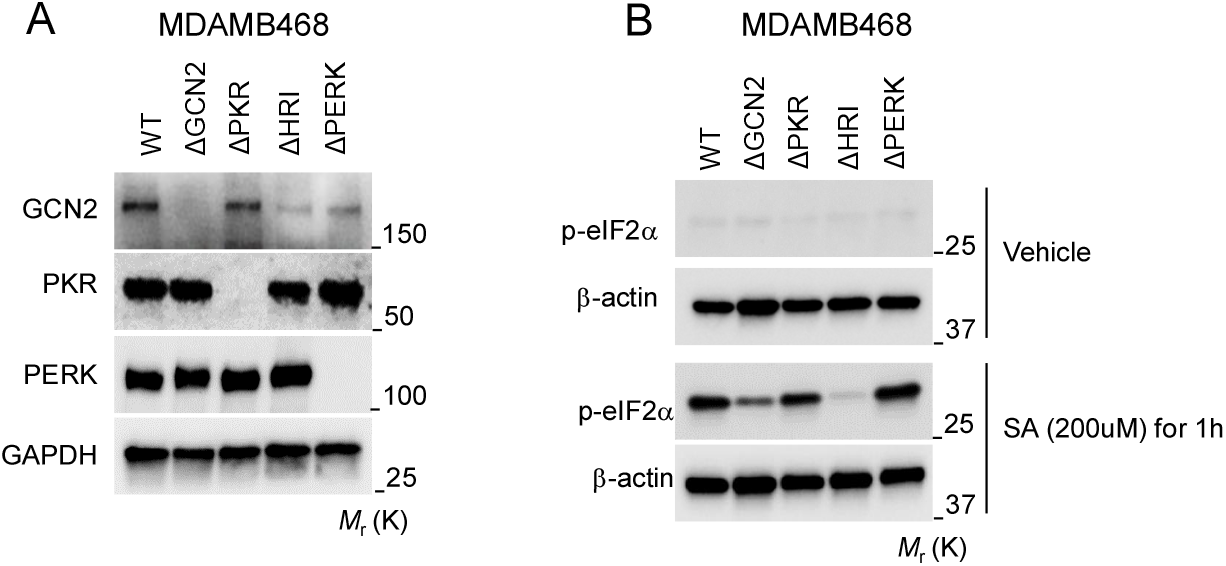
A. Immunoblot analysis of the protein levels in WT, ΔGCN2, ΔPKR, ΔHRI, and ΔPERK of MDAMB468 (note that we do not have good quality HRI antibodies). B. Immunoblot analysis of the protein levels with or without SA (200uM) for 1h in WT, ΔGCN2, ΔPKR, ΔHRI, and ΔPERK of MDAMB468. SA were treated to confirm the pattern of p-eIF2α in ΔHRI of MDAMB468 cells.

**Supplementary Figure S5, Related figure 5.**
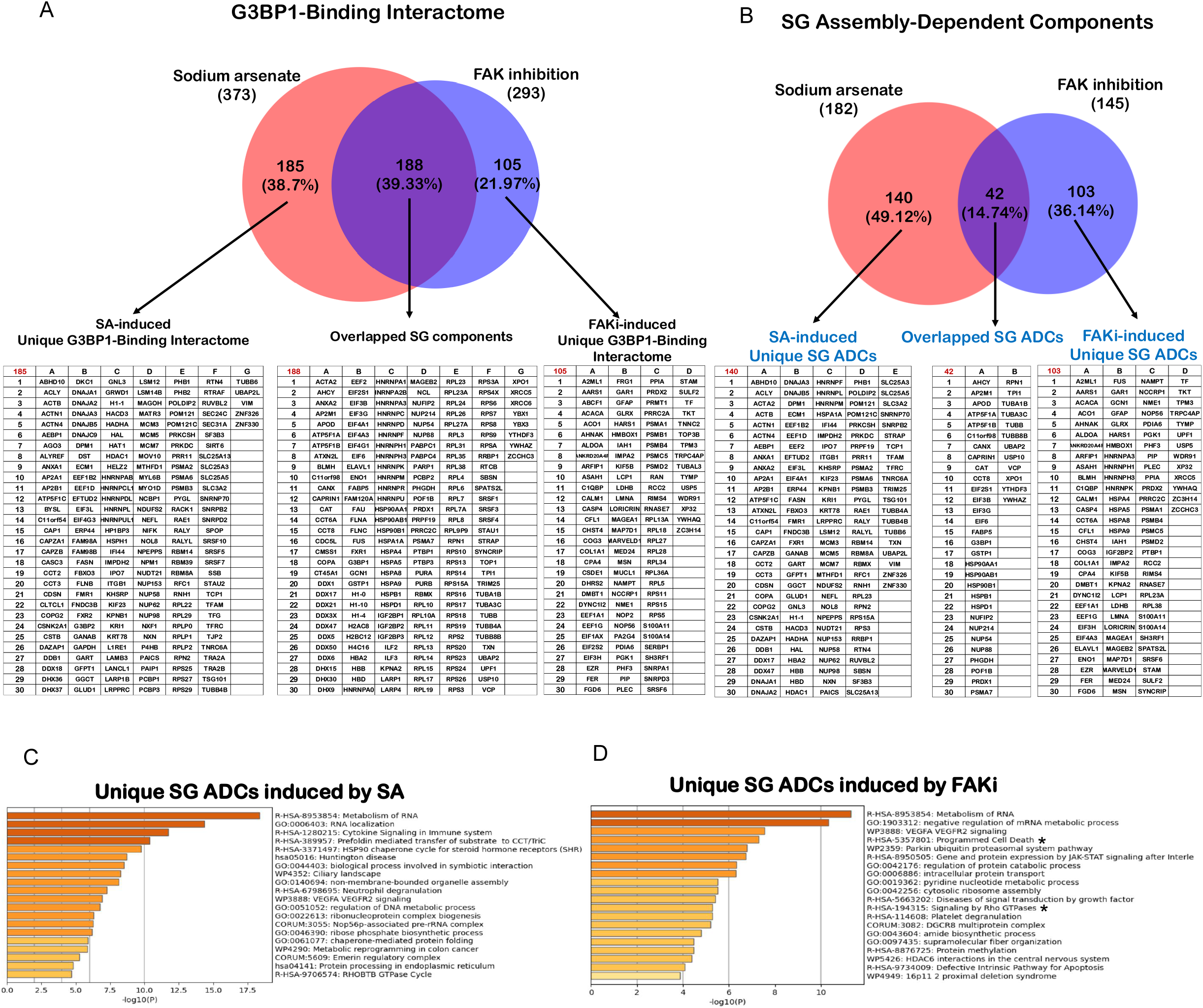
A. Identification of G3BP1-Binding Interactome in U2OS cells treated with FAKi. B. Identification of SG-ADCs in U2OS cells treated with FAKi. C. Top 20 enriched pathways up-regulated by unique SG assembly dependent component by SA. D. Top 20 enriched pathways up-regulated by unique SG assembly dependent component by FAKi.

**Supplementary Figure S6, Related figure 6.**
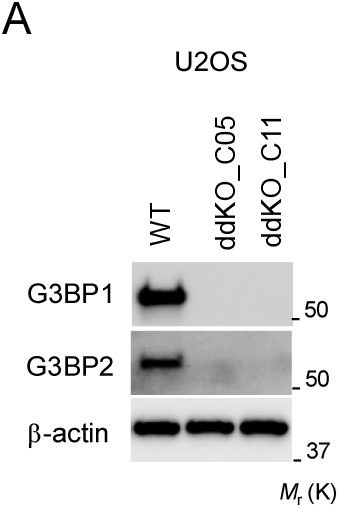
A. Immunoblot analysis of the protein levels in U2OS.

**Supplementary Figure S7, Related figure 7.**
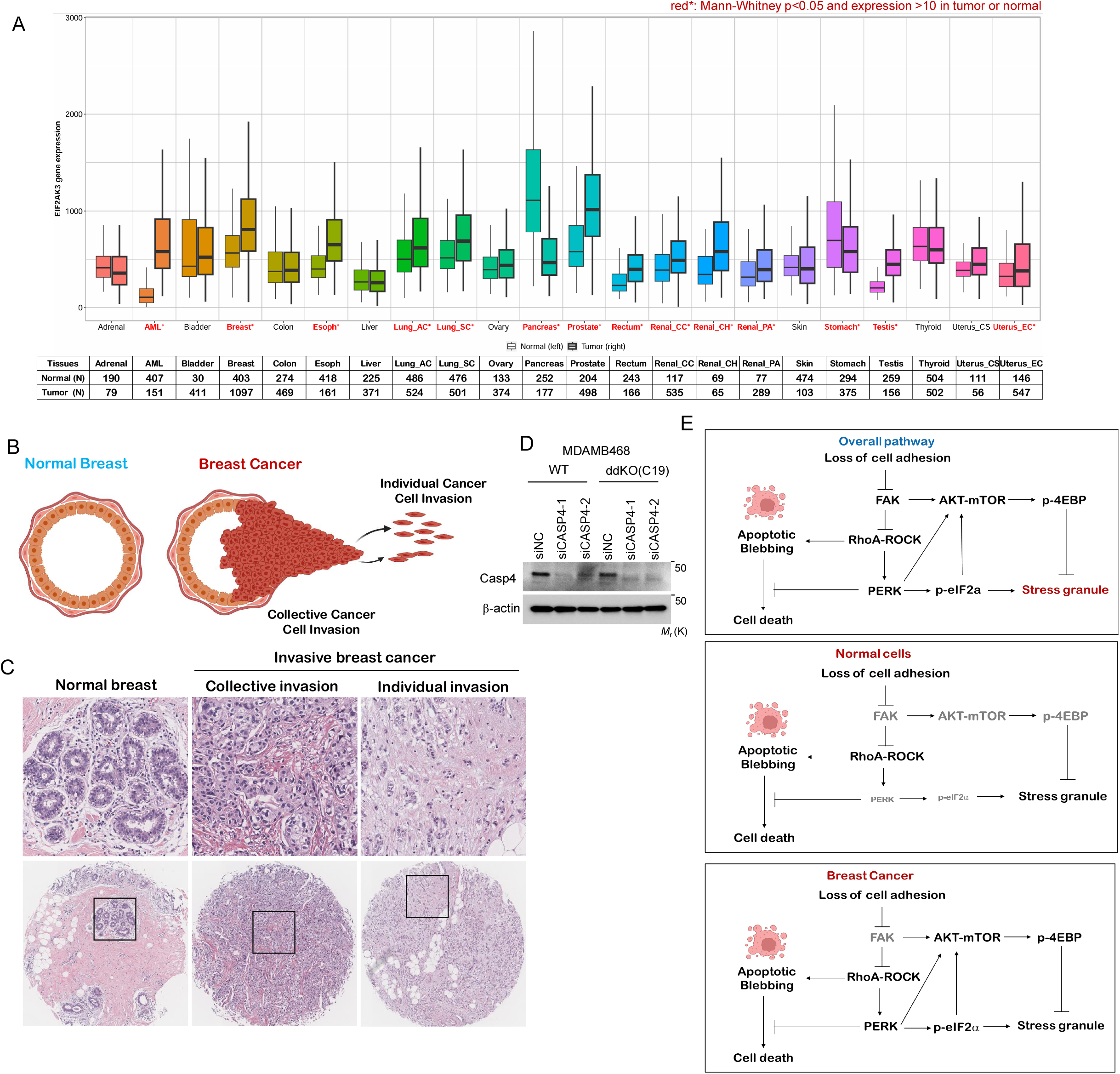
A. Differential gene expression analysis in Tumor, Normal, and Metastatic tissues (https://tnmplot.com/analysis). B. Schematic summary how breast cancer tissue was classified into normal breast, collective cancer cell invasion, or individual cancer cell invasion. C. H&E staining images provided by Tissue Mictoarray_BR087e for Fig. 7G (https://www.tissuearray.com/tissue-arrays/Breast/BR087e) D. Immunoblot analysis of the CASP4 protein levels in MDAMB468 terated with two non-overlapping siRNAs. E. A diagram illustrating the molecular mechanisms by which cells activate Stress Granules (SG) in response to weakened adhesion signals or anoikis stress to enhance anoikis resistance

